# Pan-Cancer Biomarker Analysis from the Cancer Dependency Map: A Blueprint for Precision Oncology

**DOI:** 10.1101/2025.02.07.637152

**Authors:** Dennis Gong

## Abstract

The diversity of therapeutic modalities and targets continue to grow, and the number of cancer patients eligible for targeted therapy has expanded accordingly. Despite advances, response rates to individual targeted therapy drugs remain variable, rarely achieving uniform tumor shrinkage across patients. Biomarkers are crucial for identifying patients likely to respond to therapy while sparing non-responders from toxicity and guiding them toward alternative treatments. We conducted a multi-omic analysis of biomarker-dependency relationships across 1,150 cancer molecular profiles to identify novel biomarkers for patient stratification. We first validated the Cancer Dependency Map’s sensitivity and specificity in predicting therapeutic windows for targeted therapies. Next, we identified predictive biomarkers for single- and multi-gene dependencies, assessing their selectivity within biomarker-defined populations. Protein abundance analysis revealed potential immunohistochemistry (IHC) biomarkers for clinically deployed therapeutic compounds. We performed an analysis of lineage-enriched dependencies, highlighting new opportunities for target validation and drug development. Finally, we screen for associations between hotspot mutations, damaging mutations, and protein abundance, providing insights for developing heterobifunctional small molecules for induced proximity and protein degradation. These findings advance our understanding of cancer dependencies and inform biomarker-driven strategies to optimize therapeutic outcomes.

## Introduction

Targeted therapy drug approvals are now the dominant share of new drugs for cancer patients^1^. The basic principle of these drugs is to inhibit a specific protein critical to the survival of cancer cells, but tolerable when inhibited in normal cells, thus providing a therapeutic window. These targets often belong to key growth pathways that become hyperactive in cancer due to genetic alterations. For example, gain of function mutations in growth receptor tyrosine kinases, kinase switch proteins, or the critical kinases themselves (e.g. *EGFR*, *KRAS*, *BRAF* respectively) are some of the most commonly mutated oncogene targets and are commonly therapeutically targeted^2^.

Nearly all targeted therapies are approved alongside companion diagnostic tests, which match treatments to tumors with molecular profiles most likely to respond^3^. For example, mutations identified from tumor DNA sequencing are commonly used to match patients to small molecule drugs that specifically target said mutations^4^, and IHC protein expression can be used to nominate patients for antigen targeted strategies such as chimeric antigen receptor T cells (CAR-T) or antibody drug conjugates (ADCs)^5^. Biomarkers based on RNA or protein signature scores are also emerging^6,7^, such as scores of whether a tumor is immune cold or hot^8^, or subtypes of pancreatic cancer^9^ that may have differential response to chemotherapy or targeted therapy.

Statistical frameworks have been useful for both target identification (e.g. GWAS, sequencing tumors and comparing mutations vs normal) and biomarker discovery (e.g. RNA differential expression for signatures). Statistical approaches are useful due to the large degree of molecular heterogeneity both intratumorally and across patients. The Cancer Dependency Map^10^ (DepMap) is a collection of more than 1,000 cell line models from diverse tumor types and genetic backgrounds, and have previously enabled studies of cancer dependencies and led to therapeutic targets (e.g. *WRN*^11^ dependency in cell lines with MSI high status, *DCAF5*^12^ dependency in *SMARCB1* mutant cancers). The large cohort size is a distinct advantage that enables systematic discovery and evaluation of dependency - biomarker relationships such as synthetic lethal^13^ and paralog dependencies^14^.

A variety of approaches have been implemented to predict dependency based upon a transcriptome profile from human tumors (e.g. bulk RNAseq) using DepMap as training data^15^. While tumor RNA-seq–based dependency predictions hold promise, several limitations hinder their immediate clinical utility. These challenges include stromal contamination, difficulty in distinguishing cancer-specific dependencies from normal tissue essential genes, and difficult interpretability. We propose that identifying biomarker – dependency relationships and translating these into companion diagnostic tests is a more tractable way of making translational use of the expanding use of -omic profiling technologies on patient samples.

In this study, we leveraged DepMap to identify predictive biomarkers of cancer cell dependency, including lineage-, omics-, and mutation-associated dependencies. We further focus on clinically tractable targets, which have targeting compounds in advanced clinical trials and evaluate how biomarkers can guide their use across all cancer cell lines. Additionally, we identified proteins linked to specific mutations, which may be candidates for targeted protein degradation strategies. Overall, our results provide a discovery blueprint for identifying molecularly defined patient populations and their therapeutic sensitivities.

## Results

### Sensitivity of DepMap for Detecting Therapeutic Targets

We first aimed to systematically evaluate the sensitivity of DepMap in identifying known therapeutic targets in cancer. To do so, we compiled a list of FDA-approved targeted therapies and their respective molecular targets across both solid and hematologic malignancies (n = 71). For each therapeutic target, we retrieved its corresponding Chronos dependency scores from the DepMap portal^16^. These scores provide a normalized measure of gene essentiality, where lower values indicate a greater fitness disadvantage upon knockout, while scores near zero are typically observed for non-expressed genes or those without a knockout phenotype. We hypothesized that a greater dependency (i.e., lower Chronos scores) in the approved indication compared to other cancer cell lines would provide statistical evidence of selective targeting.

We plotted several of the successfully predicted target indication pairings as density histograms representing the distribution of dependency in the target indication cell lines versus all other cell lines (**Fig. 1A**). There was variance in how closely DepMap reflected the clinically observed selectivity of therapeutic targeting. For example, in cell lines of the chronic myeloid leukemia (CML) lineage, which are commonly driven by t(9;22) BCR-ABL kinase translocations, *ABL1* was a strongly selective dependency (**Fig. 1A**), reflecting the clinical experience for patients on small molecule ABL kinase inhibitors. All but three CML cell lines met the threshold for strong dependency (ie. the average dependency of essential genes; Chronos < -1), with no cell lines outside of the CML lineage approaching strong dependency.

**Figure 1:**
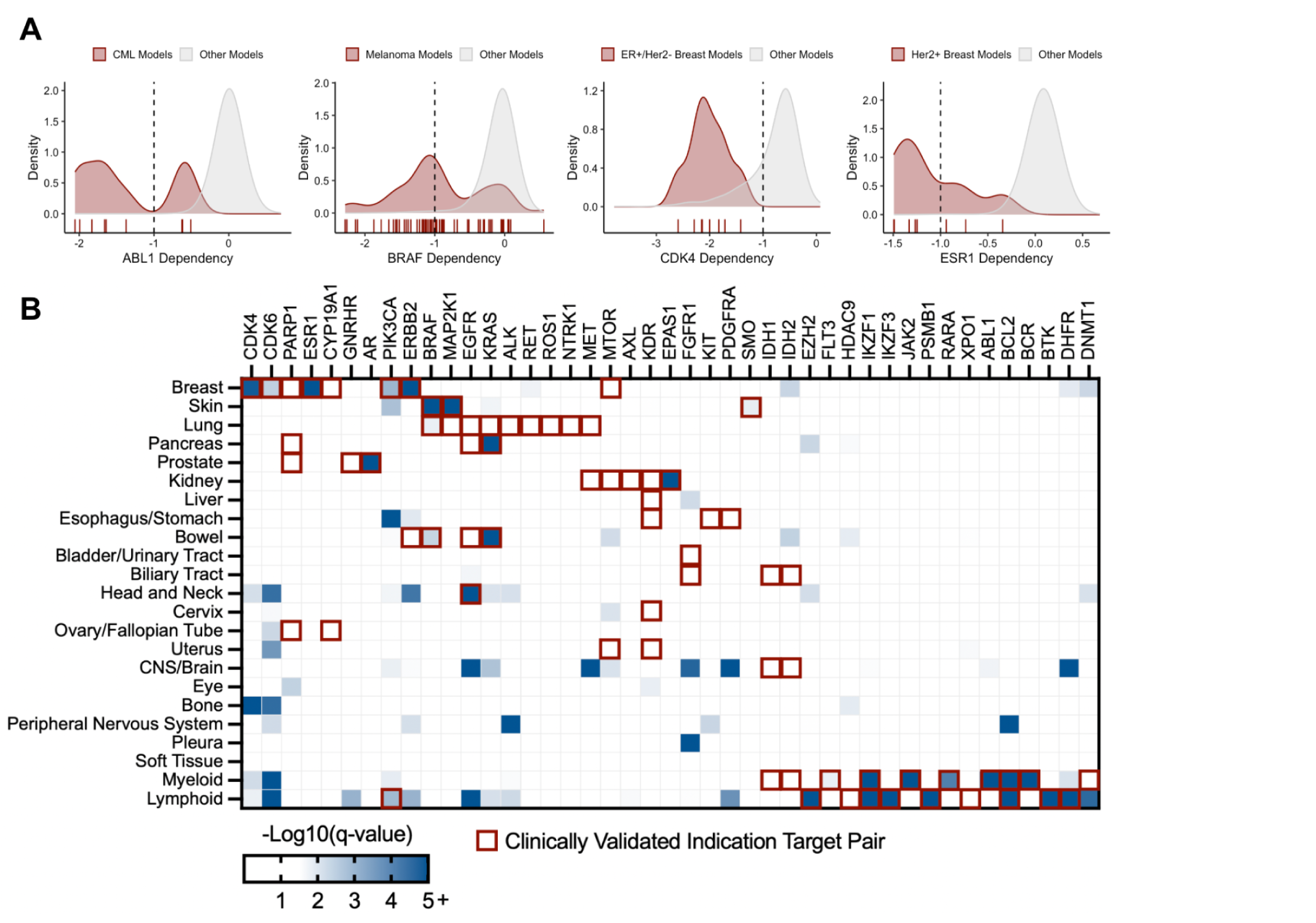
Sensitivity of DepMap for identifying anti-cancer targets. A) Example distributions of dependency scores of targets with approved therapies. stratified by the indicated clinical population versus all other cancer cell lines in the CCLE. The dotted black line at x = -1 indicates a cutoff used to define strong dependency. B) Heatmap depicting dependency (x-axis) enrichment in specific lineages (y-axis). Darker boxes indicate greater dependency in the noted lineage relative to all other lineages. Red outlines indicate manually curated clinically validated indication target pairs.

Other indication-enriched dependencies included *BRAF*, a strong dependency in 33 of 74 (44.6%) skin cancer cell lines, and *CDK4*, a strong dependency in 25 of 51 (49.0%) breast cancer cell lines and all of the ER+/Her2-lines where the drug is approved for. While these dependencies are enriched in their respective approved indications, they are also observed in cell lines from other lineages. However, both distributions exhibit significant leftward skew, such that the population means do not classify these genes as common essentials (mean Chronos score < -1). The enrichment of these dependencies within biomarker-defined or lineage-selected groups suggests a potential therapeutic window, where targeted inhibition could be selectively effective in biomarker-positive cells while sparing most other cell lines and lineages.

We systematically evaluated the indication selectivity of approved targeted therapies by conducting a t-test to compare CRISPR dependency scores between cell lines from the target molecule’s approved indication and all other cell lines (**Methods**). Enrichment was visualized in a heatmap of log-transformed adjusted p-values, and we calculated the proportion of target–indication pairs with stronger dependency in the corresponding indication lineage versus all other cell lines (**Fig. 1B**). DepMap demonstrated an overall sensitivity of 40.8% (29 validated target–indication combinations out of 71 with clinical proof of concept), significantly exceeding the pretest probability of 2.86% (12,090 significant relationships out of 422,050 possible pairings). However, the positive predictive value (PPV) of DepMap remains low (0.59%), largely due to the limited number of targets with available compounds, the evolving clinical trial landscape, and the saturation of targeted therapies in certain indications with established treatments.

DepMap performed most poorly for many solid tumor lineages including liver, lung, esophageal/stomach, bladder/urinary tract, biliary tract, cervix, ovary/fallopian tube, uterus, and CNS/brain cell line models. Overall, there was better performance in liquid tumors of either lymphoid or myeloid origin. DepMap also performed poorly for targets for which part of their mechanism of action is microenvironment mediated (e.g. angiogenesis inhibitors targeting *VEGFR2* which is encoded by *KDR*, *IDH1/2* inhibitors^17^, *GNRHR* agonists and antagonists which act on the pituitary gland, *CYP19A1*, the gene encoding for aromatase). Certain validated targets were not indication enriched because knockout is broadly toxic to all cell lines irrespective of lineage (e.g. *MTOR*, *HDAC9*, and *XPO1*). *FGFR1*, *AXL*, *KIT*, *PDGFRA* which are targets of drugs approved in tumors of the bladder/urinary tract, esophagus/stomach, kidney, and biliary tract do not show enrichment. One possible reason is that drugs targeting these proteins (e.g. imatinib, erdafitinib) have historically been multi-targeted kinase inhibitors that have pleiotropy and target multiple proteins simultaneously. Altogether removing these target indication combinations that DepMap was likely to perform poorly on (n = 25), the sensitivity improves to 63%.

There was poor performance in several lung cancer targets defined by genomic fusions (e.g. *ALK*, *RET*, *ROS1*, *NTRK1*). While these rearrangements are rare in patients^18^, there were indeed cell lines harboring these alterations, but their CRISPR knockout had negligible impact. Additionally, DepMap did not recover sensitivity to PARP inhibition or MET inhibition in any validated lineage. Interestingly, *DNMT1* dependency was strongly enriched in lymphoid cancer models despite targeting drugs (e.g. decitabine, azacitidine) only having approvals for myeloid malignancies such as AML, CMML, and MDS.

Thus, our analysis demonstrates that DepMap provides a valuable statistical framework value for identifying clinically tractable therapeutic targets.

### Predictability of Genetic Dependencies

Having established that DepMap provides significant predictive value for identifying dependencies, we moved on to systematically analyze all dependency – biomarker relationships with the goal of nominating other strong associations. The predictability of any given dependency is summarized by a random forest model trained to predict a continuous Chronos dependency score using a set of biomarkers including mRNA expression, damaging mutations, driver mutations, hotspot mutations, lineage annotation, fusion, copy number, and confounder experimental covariates. The advantage of the random forest approach is that it reduces co-linearity, nominating features that are most predictive amongst other highly correlated features. All features are also each associated with a Pearson regression coefficient (r) ranging from 0-1 quantifying the correlation of the feature with the continuous dependency score. Individual dependency – biomarker relationships are then quantified using a Gini feature importance metric (G) representing the directional contribution of the feature in the random forest model relative to the other features available in the model^19^.

In total, there are 18,443 dependencies with dependency – biomarker relationships; however, 7,511/18,443 = 40.7% of the best predictors are confounders, meaning that the effect size of the most predictive feature was less than the intrinsic screen to screen variability. We filtered out dependencies whose most predictive feature was a confounder, and also a set of poorly expressed olfactory receptor family genes whose most predictive feature was its own copy number (*n* = 101). We thus proceeded with the remaining 10,831 dependencies.

We first visualized the distribution of Pearson regression coefficients between dependency scores of each gene, and the predicted dependency. (**Fig. 2A**). The distribution was rightward skewed with a Pearson regression coefficient mean of 0.171. Roughly 2.2% of all dependencies have more than half the variance (*r* > 0.5) explainable by omics (236/10,831), which we categorized as ‘predictable’ dependencies. We also plotted the Gini feature importance (**Fig. 2B**) which we noted were also strongly right skewed. We defined ‘predictive’ features as those capable of explaining more than 10% of the variance in dependency, representing 526/10,831 (4.86%) of dependency – biomarker relationships.

**Figure 2:**
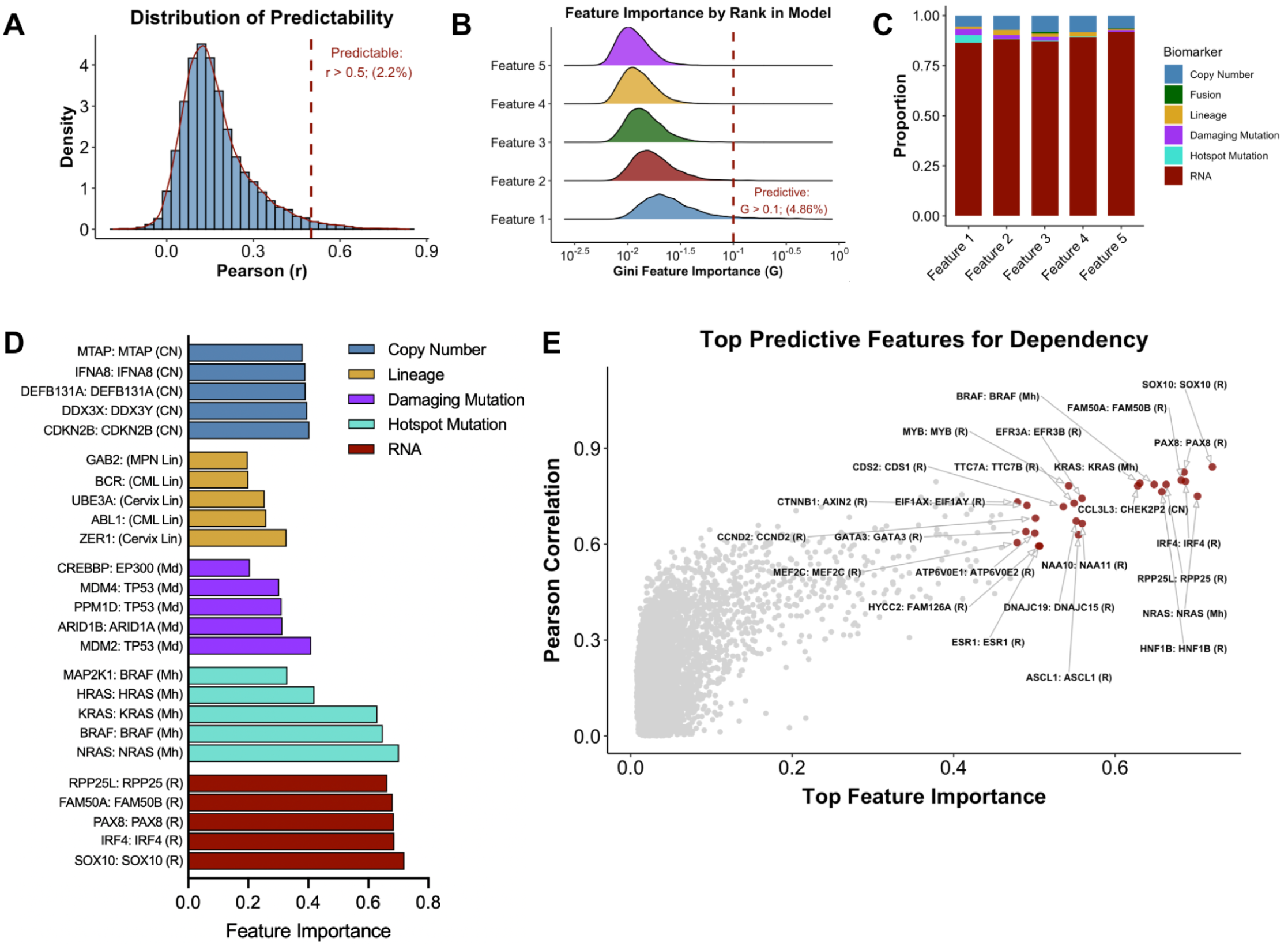
Predictability of genetic dependency. A) Histogram depicting distribution of predictability across all dependencies. Dashed line indicates cutoff used for classifying “predictable” dependencies in this paper’s analysis. B) Distribution of feature importance scores (G; x-axis) for each of the 1st through 5th top predictive biomarkers for each dependency. Dashed line indicates cutoff used for classifying “predictive” dependencies. C) Stacked barplot of the feature types represented in the 1st through 5th top predictive biomarkers for each dependency. D) Grouped barplot of top five biomarker dependency relationships in each biomarker category. The format for each label is [Dependency]:[Biomarker], where biomarker abbreviations are as follows. Md: damaging mutation; Mh: hotspot mutation; Lin: Lineage; CN: Copy number; R: RNA expression E) Scatterplot of Pearson correlation coefficients (r; y-axis) versus the most significant feature’s importance metric (G; x-axis) for all dependencies and associated biomarkers. Colored dots represent the top 25 predictive biomarkers and associated dependencies.

Gene expression has previously been reported to be the strongest predictor of in vitro cancer cell vulnerabilities^19^, and we identified the same result (**Fig. 2C**). This manifested in the form of paralogs and genes whose dependency is best predicted by its own expression. 148 of 10,831 (1.4%) relationships have a paralog or closely related member as the greatest predictor and 367 of 10,831 (3.4%) relationships have the gene itself as the greatest predictor. For example, the top biomarker relationships were often the gene expression of transcription factors with their own dependency (**Fig. S1A**).

We visualized the top five biomarker dependency relationships for each biomarker category (**Fig. 2D**). Within damaging mutations, *TP53* mutation strongly predicted *MDM2*, *MDM4*, and *PPM1D* dependency, and *ARID1A* mutation strongly predicted *ARID1B* dependency. *MDM2* and *MDM4* are negative regulators of p53 that are often overexpressed in cancer^20^, whereas *PPM1D* attenuates the stress response to DNA damage, radiation, chemicals, and chemotherapy and is activated by p53^21^. *ARID1A/B* are members of SWI/SNF protein complexes that regulate chromatin accessibility. Loss of *ARID1B* in the context of mutation of *ARID1A* has been found to destabilize SWI/SNF and impair proliferation^22^.

Highly predictive hotspot mutations were saturated with oncogenes (*BRAF*, *HRAS*, *NRAS*, *KRAS*) (**Fig. S1B**). Mutations in these proteins have been used as patient selection biomarkers for oncogene directed therapies^23^. The top lineage predictive dependency was *ZER1* predicted by cervical cancer lineage, which was notable due to the molecular role of *ZER1* in helping HPV E7 protein^24^, and the clinical association between HPV and cervical cancer. Within copy number biomarkers, the *DDX3X* and *DDX3Y* paralog pair^25^ stood out as another paralog pair with potential for targeting given the significant dependency on *DDX3X* with *DDX3Y* loss. Loss of the Y chromosome, where *DDX3Y* is located, is clonal^26^ across several cancers in males. Finally, the most predictive category of gene expression was abundant in both paralog pairs (e.g. *EPP25L* dependency predicted by *RPP25* expression) and dependencies correlated with self expression. In particular, transcription factors *PAX8*, *IRF4*, *HNF1B*, and *SOX10* **(Fig. S1A**) were amongst the most predictable dependencies across any biomarker dependency pairing.

Finally, we visualized all biomarker dependency relationships on a scatterplot to highlight the strongest biomarker relationships (**Fig. 2E**). Reassuringly, several biomarker defined therapeutic strategies in clinical use (e.g. BRAF targeting agents in *BRAF* mutant defined populations, trials of KRAS inhibitors in *KRAS* mutant patients, hormone therapy targeting *ESR1* in ER+ breast cancer patients). Thus, our analysis defines a landscape of new opportunities for biomarker directed therapy.

### Biomarkers of Clinically Tractable and Selective Dependencies

We identified clinically tractable dependencies by querying OpenTargets^27^, a publicly available resource compiling clinical trials and drug molecules targeting therapeutic targets. We defined clinically tractable targets as those with associated targeting molecules (e.g. small molecule inhibitors, monoclonal antibodies, small molecule degraders) with Phase I clinical trial data (**Methods**).

We plotted these targets on a scatter plot overlaying a selectivity metric (x-axis) and their predictability (y-axis) reasoning that the selectivity axis may nominate targets with enhanced therapeutic window, and the predictability axis may nominate targets with readily available biomarkers (**Fig. 3A**). While there were a few targets that were both selective and predictable (e.g. *KRAS*, *BRAF*, *NRAS*, *CTNNB1*, *ERBB2*, *FGFR1*, *MET*), the majority of clinically tractable targets (76.5%) were neither predictable (predictability (*r*) <= 0.5) nor selective (defined as CRISPR LRT score <= 100) (**Methods**). However, 189/233 = 81.1% of predictive dependencies were selective (**Fig. 3B**), highlighting the fact that biomarker predicted dependencies are likely to have a selective therapeutic window.

**Figure 3:**
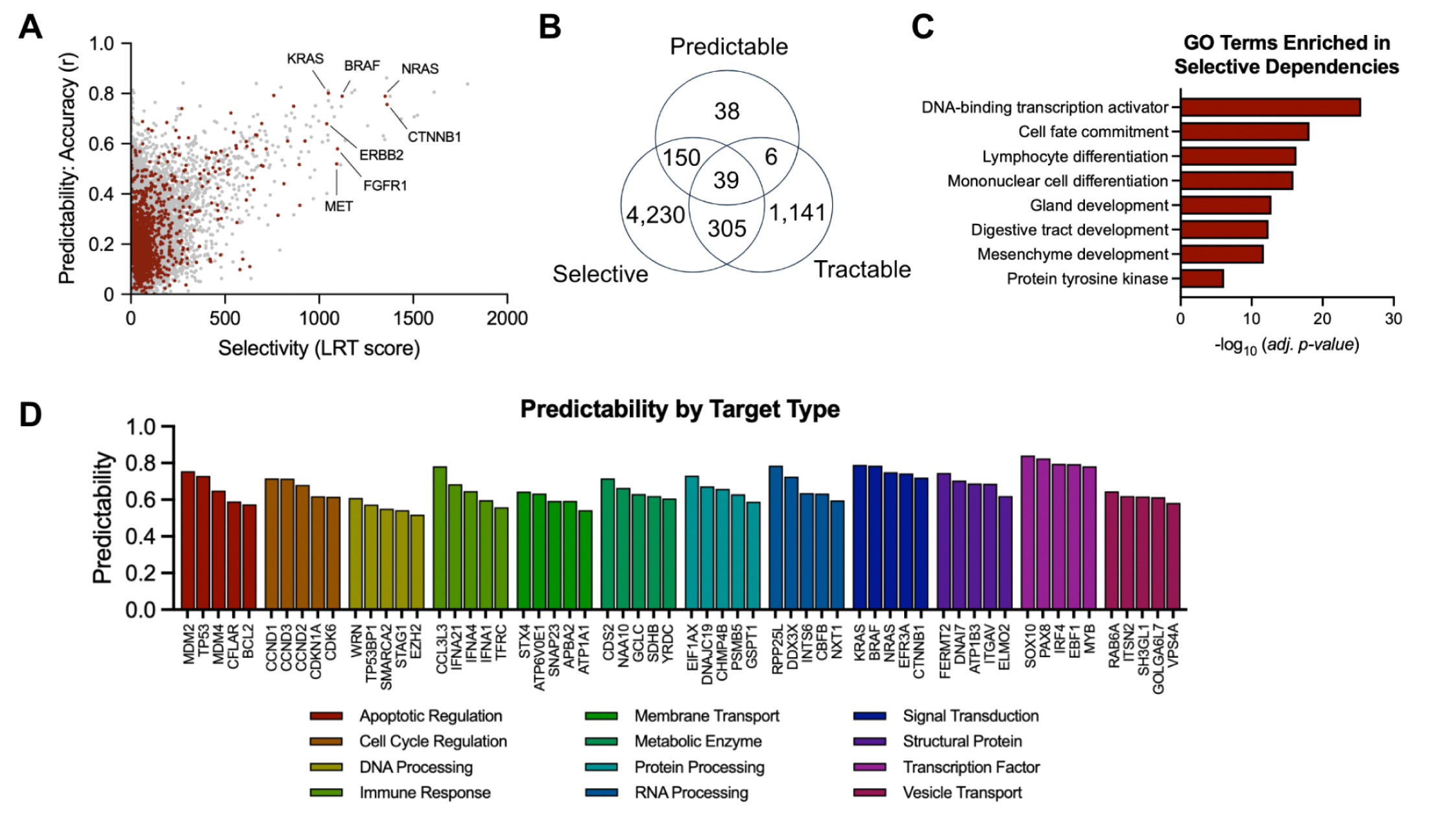
Biomarkers of clinically tractable and selective dependencies. A) Scatterplot of selectivity versus predictability. Clinically tractable targets are highlighted (red dots). B) Venn diagram depicting overlap of selective, clinically tractable, and predictable dependencies. C) Barplot of top enriched GO terms from GSEA analysis of selective dependencies (ie. LRT score > 100). D) Barplot of selective dependencies grouped by their protein class and their predictability (y-axis). The top five predictable dependencies for each protein class are visualized.

Selective dependencies were enriched for protein kinases, but most strongly by lineage defining gene sets including specific terms for mesenchymal, digestive tract, gland, and lymphoid differentiation and development (**Fig. 3C**). Activating transcription factors were the primary protein class to be enriched in selective dependencies (**Fig. 3C**). We classified the other selective targets into their functional protein classes and plotted the top 5 dependencies in each target class ranked by their predictability by biomarkers (**Fig. 3D**). We found that expectedly, transcription factors are most readily predicted, followed by signal transduction pathway components (**Fig. 3D**). However, a diverse set of protein classes were represented including apoptotic regulatory proteins, cell cycling and DNA replication proteins, metabolic enzymes, RNA/protein processing enzymes, signal transduction components, and structural proteins, which reflect the breadth of mechanisms currently used as cancer therapy.

During our review of ‘predictable’ dependencies (ie. random forest model Pearson *r* > 0.5), we noticed that several genes were biomarkers of multiple dependencies. These were less common, representing just 56 unique biomarkers, but they altogether made up 14% of ‘predictable‘ dependency – biomarker relationships (**Fig. 4A**). Most commonly, a biomarker was predictive of two dependencies. The most common biomarker of dependency was the gap junction gene *GJA1* which encodes for connexin 43 and is associated with reduced dependency to 20 distinct genes (**Fig. 4A**). These 20 genes were all related to metabolic function, playing critical roles as enzymes (e.g. *SDHB*, *YRDC*, *ADSL*) or other nutrient transporters (e.g. *TRPM7*). These genes tended to also to be essential genes (Chronos < -0.5), such that knockout often had deleterious effects. Thus, we hypothesized that cells with enhanced *GJA1* expression may enable connection with neighboring cells and nutrient sharing, thus compensating for genetic knockout of the critical metabolic factor (**Fig. 4B,C**).

**Figure 4:**
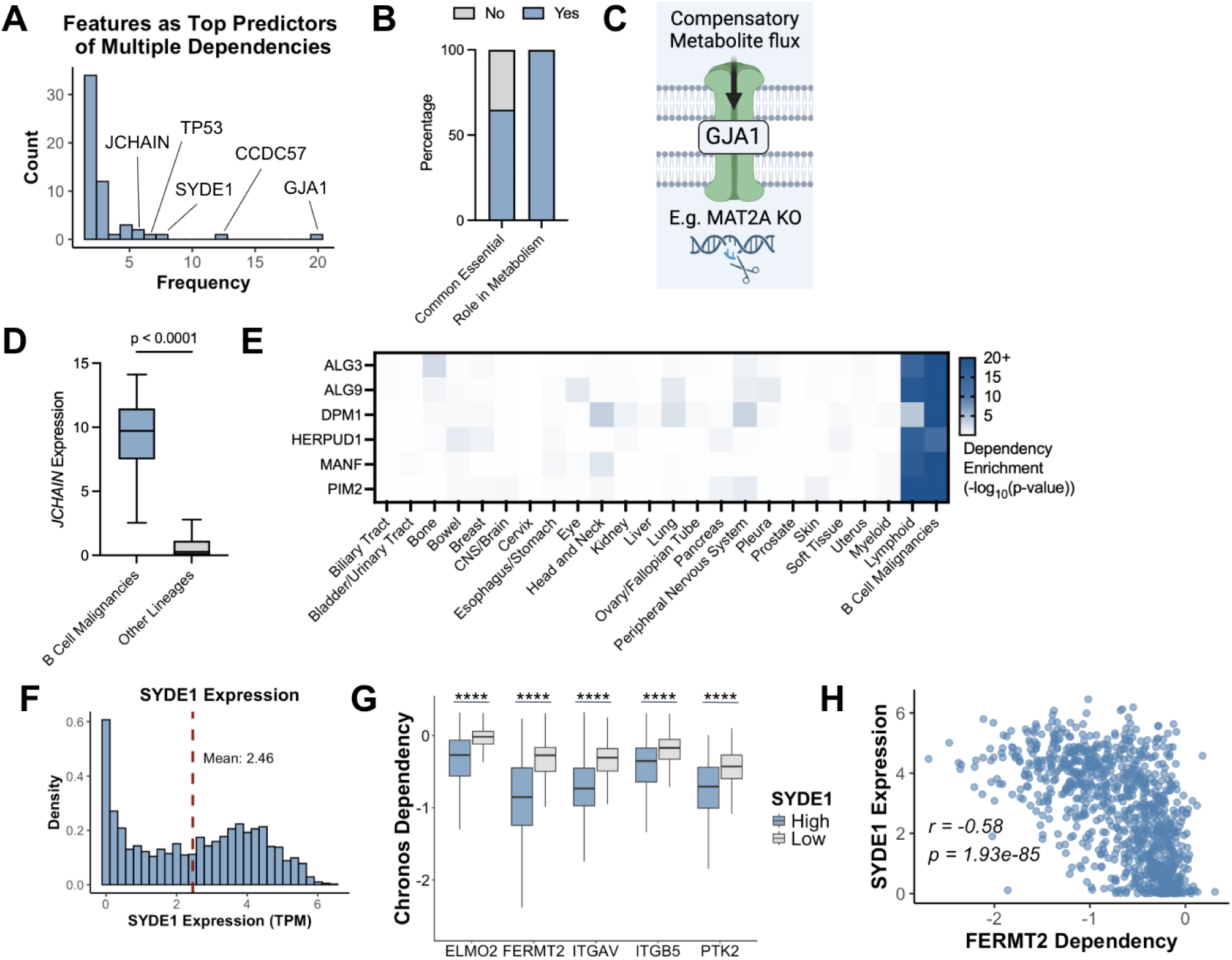
Biomarkers of multiple dependencies. A) Histogram showing distribution of predictors of more than 1 dependency. B) Proportion of *GJA1* predictable dependencies that have Chronos dependency score < -0.5 and have roles in metabolism. C) Theoretical model for how *GJA1* expression reduces dependency on critical metabolic enzymes. D) Boxplot showing *JCHAIN* expression in B cell malignancies versus all other cell lines. E) Heatmap showing dependency enrichments of *JCHAIN* predictable dependencies in each lineage. F) Histogram of *SYDE1* expression across all profiled cancer cell lines. Red dashed line indicates mean *SYDE1* expression and cutoff used to define *SYDE1* high vs low cells. G) Grouped boxplots for dependencies predictable by *SYDE1* expression. Comparisons are made between cell lines above and below the mean *SYDE1* expression across all profiled cancer cell lines. **** indicates p < 0.0001. H) Scatterplot of *SYDE1* RNA expression (y-axis) versus FERMT2 dependency (x-axis).

Another common multi-dependency biomarker finding was the association of lineage specific dependencies with lineage specific transcripts. The most notable example was B cell associated transcripts (e.g. *JCHAIN* RNA expression) and B cell malignancy dependencies such as the mannosyl transferases *ALG3*, *ALG9*, and *DPM1*, the serine/threonine kinase *PIM2*, and the ER stress proteins *HERPUD1* and *MANF* (**Fig. 4D,E**).

We found that the Rho GTPase-activating protein encoding gene *SYDE1* was a predictor of multiple dependencies associated with focal adhesion signaling. Using a cutoff of mean *SYDE1* expression across all solid tumor cell lines, we defined *SYDE1* high and low groups (**Fig. 4F**) that significantly stratified dependency to focal adhesion pathway proteins *ELMO2* (*p* = 8.28e-61), *FERMT2* (*p* = 7.02e-71), *ITGAV* (*p* = 1.50e-62), *ITGB5* (*p* = 2.49e-27), and *PTK2* (*p* = 3.35e-41) (**Fig. 4G,H**). Inhibition and degradation of the focal adhesion kinase (FAK) encoded by *PTK2* is currently an actionable strategy for certain MAPK driven cancers but is limited by heterogeneous single agent activity^28^. Thus, further investigation of *SYDE1* expression as a biomarker for FAK inhibition sensitivity is warranted and may have clinical utility. Other recurrent biomarker dependency relationships of potential clinical significance included *TP53* mutation as a biomarker of *TP53* degradation and regulator dependencies including *MDM2, MDM4, PPM1D, CHEK2, UBE2D3, DDX31, and USP7*. Additionally, *BRAF* mutation predicts MAPK pathway members *SHP2*, *ERK2*, *MEK1*, and *BRAF* itself.

Finally, we sought to assess the importance of our clinically tractable, selective, and predictable dependencies across each of the individual cell lines contained within DepMap. Our aim was to survey the landscape of targets for individual cell lines, given that it has previously been said that a relatively small sliver of patients are eligible and benefiting from precision oncology medicines^29^. The majority of precision medicine strategies are based on a genetic biomarker, most commonly hotspot mutations in oncogene drivers. Thus, we wondered whether we could identify additional targets for patients using additional biomarker classes.

We began by identifying the most preferentially dependent targets for each DepMap cell line by standardizing dependency scores (Z-scores) and selecting the gene with the lowest Z-score per cell line. Our rationale was that the most preferentially dependent gene would offer the broadest therapeutic window against all other cell lines. Initially, we focused on clinically tractable and predictable dependencies before expanding our analysis to include selective and predictable dependencies. Among clinically tractable and predictable dependencies, the most preferentially dependent genes were often strong dependencies (75.2% had a Chronos score < -1; **Fig. S2A**). Across all targets showing preferential dependency (Chronos score 1.5 standard deviations below the mean) in a given cell line, 909 (79%) had at least one target with a dependency score < -1, with an average of 3.0 targets per cell line (range: 1–13). Including selective targets, 1,086 (94.4%) of cell lines had at least one strong dependency, with an average of 7.7 targets per cell line (range: 1–41) (**Fig. S2B**). The most targetable lineages included skin, breast, bowel, and liquid tumors (myeloid and lymphoid), while the least targetable were cervix, soft tissue, uterus, pleura, and thyroid (**Fig. S2C**).

The preferential targets varied widely across cell lines, underscoring the necessity of biomarkers for patient stratification. Among clinically tractable and predictable targets, each target was preferentially dependent in a mean of 44.5 cell lines (range: 1–131); this increased to 62.6 cell lines (range: 3–131) when including selective targets (**Fig. S2D**). As expected, gene expression was the strongest biomarker for most preferential and strong targets, followed by hotspot and damaging mutations (**Fig. S2E**). Using paired PRISM small-molecule screening and protein expression data, we found that gene expression not only predicts dependency but also therapeutic sensitivity to targeted compounds. The strongest predictive signal was observed for MAPK pathway dependencies, including *EGFR*, *ERBB2*, *BRAF*, *MAP2K1*, and *ERBB4* (**Fig. S3**).

Thus, we have demonstrated that DepMap is capable of identifying biomarkers that stratify dependency to selective and clinically tractable targets.

### Lineage Enriched Dependencies

Next, we systematically analyzed dependencies enriched in tissue lineages. We reasoned that this approach may nominate new biologically relevant pathways or genes that are specific to a tissue lineage, which can eventually be evaluated for their potential for therapeutic targeting. We first calculated, using our previous t-test approach, enrichment scores for each lineage relative to all other lineages, across all target encoding genes (**Fig. 5A**). Overall, we noted heterogeneity in the number of selective dependencies for each lineage but nonetheless a significant number of lineage enriched dependencies for each lineage. A majority of highlighted targets are not yet clinically tractable (12.7%), but almost all are selective (83.9%), highlighting a significant opportunity for new drug development.

**Figure 5:**
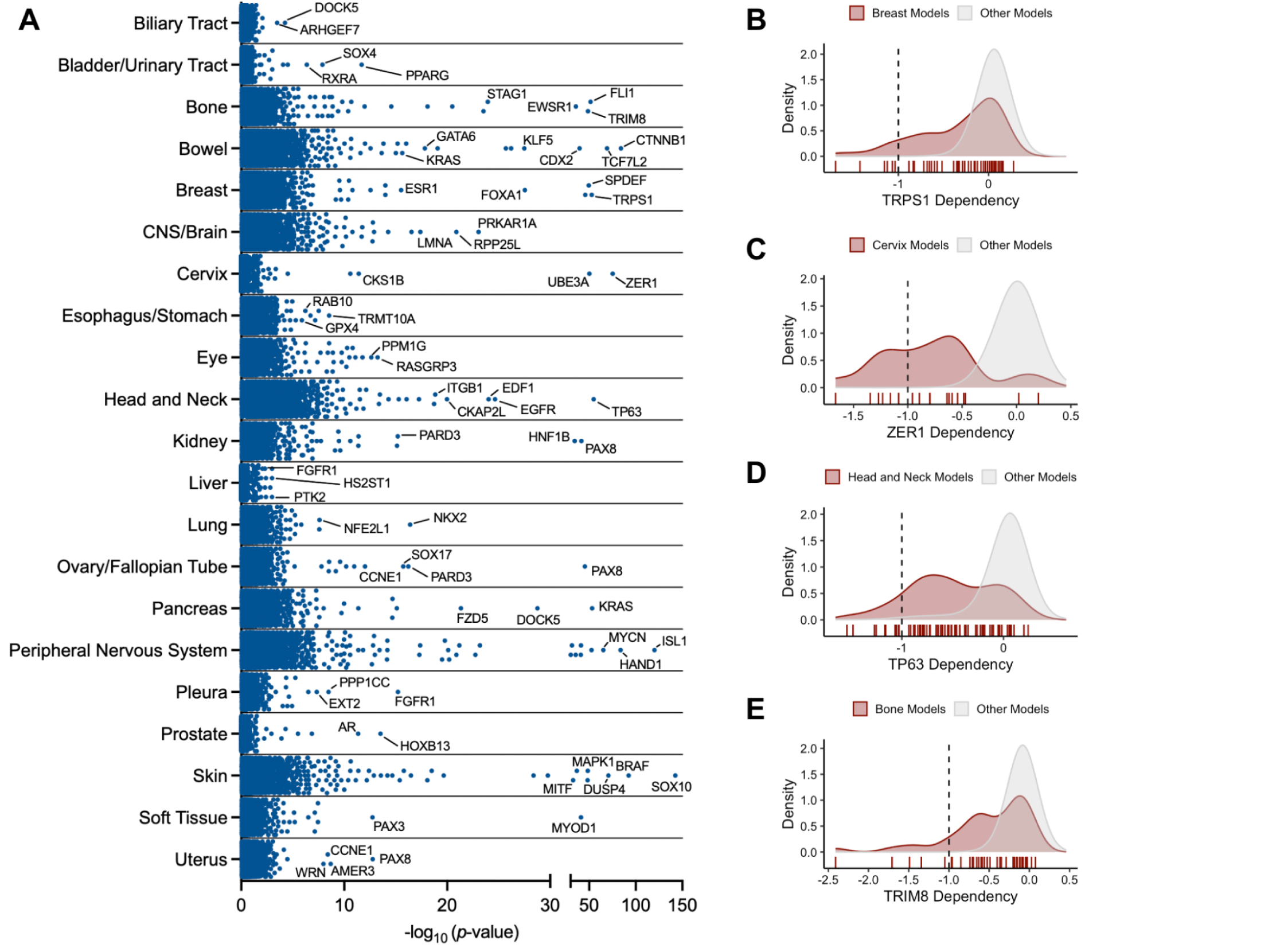
Lineage enriched and biomarker prioritized dependencies. A) Enrichment plot depicting lineage enriched dependencies, quantified by -log10(q-value) of a t test comparing dependency in stated lineage cell lines versus all other cancer cell lines. B-E) Stacked histograms showing dependency distributions in lineage of enrichment versus all other lineages for *TRPS1* in breast cancer cell lines, *ZER1* in cervical cancer, *TP63* in head and neck cancers, and *TRIM8* in bone cancers.

There were dependency similarities within lineages of similar functional roles and anatomic locations. For example, *TP63*, a master transcription factor for squamous identity, was strongly enriched in head and neck cancer cell lines, which are often squamous cell carcinomas (**Fig. 5B**). Head and neck cancers were also enriched for the endothelial differentiation factor *EDF1*, and the epidermal growth factor receptor *EGFR*. Squamous esophageal and esophagogastric cancers (i.e. esophageal squamous cell carcinoma) also had enrichment for mesenchymal dependencies such as *GPX4*^30,31^. Additional high ranking dependencies included *RAB10*, which is a small GTPase protein that has been shown to participate in the progression of breast, pancreatic, liver, and lung cancers, and *TRMT10A* which encodes a tRNA methyltransferase homolog gene.

The kidney was strongly enriched for developmental transcription factors *PAX8* and *HNF1B*. Interestingly, the kidney shared dependency on *PAX8* and developmental scaffolding protein *PARD3* with ovarian, fallopian tube, and uterine cancer cell lines, suggesting a common developmental lineage dependency in cancers arising from intermediate mesoderm tissues. In uterine, ovarian, and fallopian tube cancer cell lines, we also observed common dependency on *CCNE1*, which encodes a critical cell cycle protein. Molecular glue degraders of *CCNE1* are in development for *CCNE1* amplified tumors, which become prevalent as a resistance mechanism to CDK4/6 inhibitors^32^. Our results provide rationale for indication expansion of these drugs into reproductive system tumors. Cervical cancer cell lines were a notable exception to the shared patterns of dependency in uterine, ovarian, and fallopian tube cancer cell lines. In the cervix, enriched dependencies were involved in the stability of viral proteins including *ZER1*^24^ (**Fig. 5C**) and *UBE3A*^33^. This reflects the clinical experience of HPV infections being a causative factor for the development of cervical cancer.

In other reproductive organs, dependency on hormonal growth signals was another notable pattern. *SPDEF* and *FOXA1* are transcription factors that play roles in regulating estrogen and androgen receptor signaling, and are selectively dependent in invasive breast carcinomas and prostate carcinomas. Breast cancer cell lines were strongly dependent on the estrogen receptor gene *ESR1,* which drives hormonal growth signaling and is targetable with selective estrogen receptor degraders (SERDs). Breast cancer lines were also enriched for *TRPS1* (**Fig. 5D**), which has a role in estrogen receptor signaling and is commonly used as a diagnostic marker for breast cancer^34^. In the prostate cancer cell lines, the androgen receptor *AR* and the androgen receptor repressor *HOXB13* were enriched, highlighting the strong role of hormone signaling. *HOXB13* is an example of a gene thought to be a tumor suppressor, as hereditary mutations significantly increase the risk of prostate cancer^35^; however, acute ablation has deleterious effects on cancer cells. Pharmacological castration is also a highly effective strategy in prostate cancer, suggesting that a goldilocks level of androgen signaling is necessary for prostate cancer viability^36^.

Signaling addiction and consequent genetic dependency was another recurrent theme. In addition to hormone signaling, a broad set of other pathways were represented, including Wnt/β-catenin, PPAR, MAPK, and FGFR. In bowel (colon) cancer cell lines, we observed enrichment of *TCF7L2* and *CTNNB1*, which together form a bipartite transcription factor that enforces constitutive Wnt signaling^37^, as well as enriched dependency on the nuclear protein *CDX2*. *CDX2* is often lost due to mutation or copy number loss in colon cancer and similar to *HOXB13,* was thought of as a tumor suppressor, given that loss of *CDX2* is associated with poor prognosis^38^. *CDX2* suppresses Wnt/beta-catenin signaling^39^, suggesting that acute loss may cause deleterious overactivation of beta-catenin signaling, providing another example of the fitness advantage of a goldilocks intermediate level of signaling.

*KRAS* was the dominant pancreas cancer enriched dependency, in line with the high prevalence of KRAS gain of function mutations^40^ and the efficacy of therapeutic targeting in preclinical and clinical settings^41–43^. Pancreatic cancer cell lines also had enriched dependency for the guanine nucleotide exchange factors genes *ARHGEF7* and *DOCK5*, which were also the top two enriched dependencies in biliary tract cell lines. While there were few oncogene dependencies observed for biliary tract cell lines compared to pancreatic cancer lines, a recently published biliary tract cancer cell line atlas^44^ detected strong dependency for *KRAS, CTNNB1, BRAF, FGFR2, and EGFR*, indicating that cell line availability was a major limiting factor for dependency discovery and that pancreatic and biliary tract cancers do indeed share several core dependencies in oncogene drivers. The skin also contained strong enrichment for oncogene driver dependencies, including the therapeutic targets (*BRAF*, *MAPK1*). Notably, the loss of *DUSP4*, a MAPK suppressor, in MAPK driven skin cancers also leads to hyperactivation and cell death in these cancer cells^45^, providing a third example of the importance of goldilocks signaling. Finally, lineage-defining transcription factors including *MITF* and *SOX10* are strongly dependent in skin cancer cell lines.

In bladder/urinary tract cell lines, the nuclear receptors *RXRA* and *PPARG*, as well as the transcription factor *SOX4* were enriched. *RXRA* regulates transcription as part of a heterodimer with 14 other nuclear receptors, including the peroxisome proliferator-activated receptors (PPARs). Hyperactive PPAR signaling, either due to *PPARG* gene amplification or RXRA hot-spot mutation (S427F/Y) drives 20–25% of human bladder cancers^46^. Meanwhile, *SOX4* is overexpressed in 24% of bladder cancers but the precise role and reason for dependency is not yet elucidated. Several eye-enriched dependencies were notable including *PPM1G* and *RASGRP3* in uveal melanomas. *RASGRP3* encodes a RAS activator protein that is significantly and selectively overexpressed in response to *GNAQ*/11 mutation in uveal melanoma^47^, of which the prevalence is 80%.

*PPM1G* encodes a protein phosphatase whose role has been explored in hepatocellular carcinoma as a regulator of the splicing factor *SRSF3*, but whose role has not yet been defined in eye cancers. Pleura (ie. mesothelioma) has enriched dependency for *EXT2*, *PPP1CC*, and *FGFR1*. *FGFR1* has previously been identified as a driver of mesothelioma^48^ but selective small molecule inhibition of FGFR1,2, and 3 did not show efficacy in clinical trials^49^. The *EXT2* gene encodes a glycosyltransferase involved in synthesizing heparan sulfate proteoglycans. *PPP1CC* encodes the gamma isozyme of the PP1 subfamily of protein phosphatases. *EXT2* and *PPP1CC* both have not received significant characterization in mesothelioma.

Aberrant transcription factor utilization and genetic dependency were previously highlighted in kidney and other lineages but are also particularly apparent in soft tissue sarcomas. These cancers often have genomic rearrangements that support the overutilization of lineage-defining transcription factors *MYOD1* and *PAX3*. Overexpression of *MYOD1* in alveolar and embryonal rhabdomyosarcoma via gene amplification is critical to cancer progression, and consequently, knockout of *MYOD1* is lethal. Dependency enrichment of *PAX3* is due to the high prevalence of the PAX3-FOXO1 fusion gene in alveolar rhabdomyosarcomas^50^. In bone cancer models (Ewing sarcoma), the FLI1/EWSR1 oncogene fusion leads to constitutive activation of *FLI1* signaling and is negatively regulated by *TRIM8*^51^ (**Fig. 5E**), both of which are enriched dependencies. Ewing sarcoma models also have enriched dependency of *STAG1* due to frequent loss or mutation of its paralog pair *STAG2*^52^.

Finally, both central and peripheral nervous system cancer cell lines have unique dependency profiles relative to other solid tumors. In brain tumor models, *PRKAR1A*, *RPP25L*, and *LMNA* have enriched dependency. *PRKAR1A* encodes for protein kinase A (PKA). *RPP25* is hypermethylated in GBM^53^ and encodes a tRNA processing complex subunit, with *RPP25L* as its paralog. *LMNA* encodes for part of the nuclear lamina. Nerve sheath tumors and neuroblastomas, which often affect children, had an abundance of selective dependencies including *ISL2,* a motor neuron developmental transcription factor, and *HAND1*, a developmental transcription factor involved in cardiac muscle development, trophoblast differentiation, and yolk sac vasculogenesis.

Several lineages did not have enriched dependencies, including liver and lung cancer cell lines. This was surprising, given the frequent usage of targeted therapy in each indication (e.g. multi-kinase inhibitors in HCC and genomic alteration targeted inhibitors in NSCLC). In lung cancer cell lines, there were several enriched dependencies specific to small-cell lung cancer including *NKX2*, a highly expressed dual lung and neural lineage transcription factor, and *NFE2L1*, a transcription factor that controls the expression of stress response and cytoprotective genes in neurons. Non-small cell lung cancer cell lines were strongly enriched for *SMARCA2* dependency, a gene that encodes a subunit of SWI/SNF complexes. The liver had the fewest and weakest lineage-enriched dependencies, including *PTK2*, which encodes the central focal adhesion complex protein FAK, *FGFR1* which encodes fibroblast growth factor receptor, and *HS2ST1* which encodes a heparan sulfate sulfotransferase and is associated with worse overall survival in patients with hepatocellular carcinoma^54^.

Overall, we have defined the landscape of lineage-enriched dependencies, highlighting remarkable opportunities for new target validation and drug development.

### Protein Expression Correlated with Mutational Status

While targeted therapy relies on sensitivity to target inhibition, a new class of therapeutic modality relies on therapeutic targeting via differential protein abundance in cancer cells relative to normal cells. While differential protein abundance can be assessed in a straightforward manner through protein staining (e.g. via IHC), it has recently been reported that mutational status can also be a reliable indicator of protein abundance of certain disease associated proteins like p53^55^. Novel heterobifunctional modalities^56–58^ (e.g. Halda Therapeutics, Shenandoah Therapeutics) capable of exploiting dysregulated protein expression may benefit from novel biomarkers. Thus, we devised a statistical enrichment strategy to assess the enrichment of protein expression in molecularly defined cancer subsets.

We restricted our analysis to solid tumors given the significant differences in mutation landscape and protein expression between solid and liquid tumors. Protein expression for 214 targets are collected for approximately 84.8% (n=898) of the cell lines profiled in DepMap. For each gene mutated at least 10 times across all DepMap cell lines, we performed a t-test comparing the protein expression of each target across mutated or wild-type cell lines and calculated the Log2 fold change in expression.

We plotted volcano plots of mutation - protein expression relationships for both damaging and hotspot mutations (**Fig. 6A,B**). For both hotspot and damaging mutations, we recovered p53 mutation as a significant outlier in terms of accumulation of the mutated protein^55^ (**Fig. 6B,C**). *P53* damaging mutations also were associated with increased protein abundance of CHK2 pT68 and p14ArfBetA300-340A and decreased abundance of the pro-apoptotic protein Bax. Unlike p53, mutations in tumor suppressors *PTEN* and *RB1* significantly decreased levels of their encoded protein. *PTEN* loss of function mutations were associated with significantly higher levels of Akt phosphorylation at sites S472 and T308 (**Fig. 6B,C**). Gain of function *KRAS* mutations significantly increased *VEGFR2* protein expression (**Fig. 6C**), consistent with the angiogenic role of oncogenic *KRAS* activation. *BRAF* hotspot mutations were associated with increased *MEK1* phosphorylation at S217 and S221 but decreased C-RAF pS338 and SHP2 pY542.

**Figure 6:**
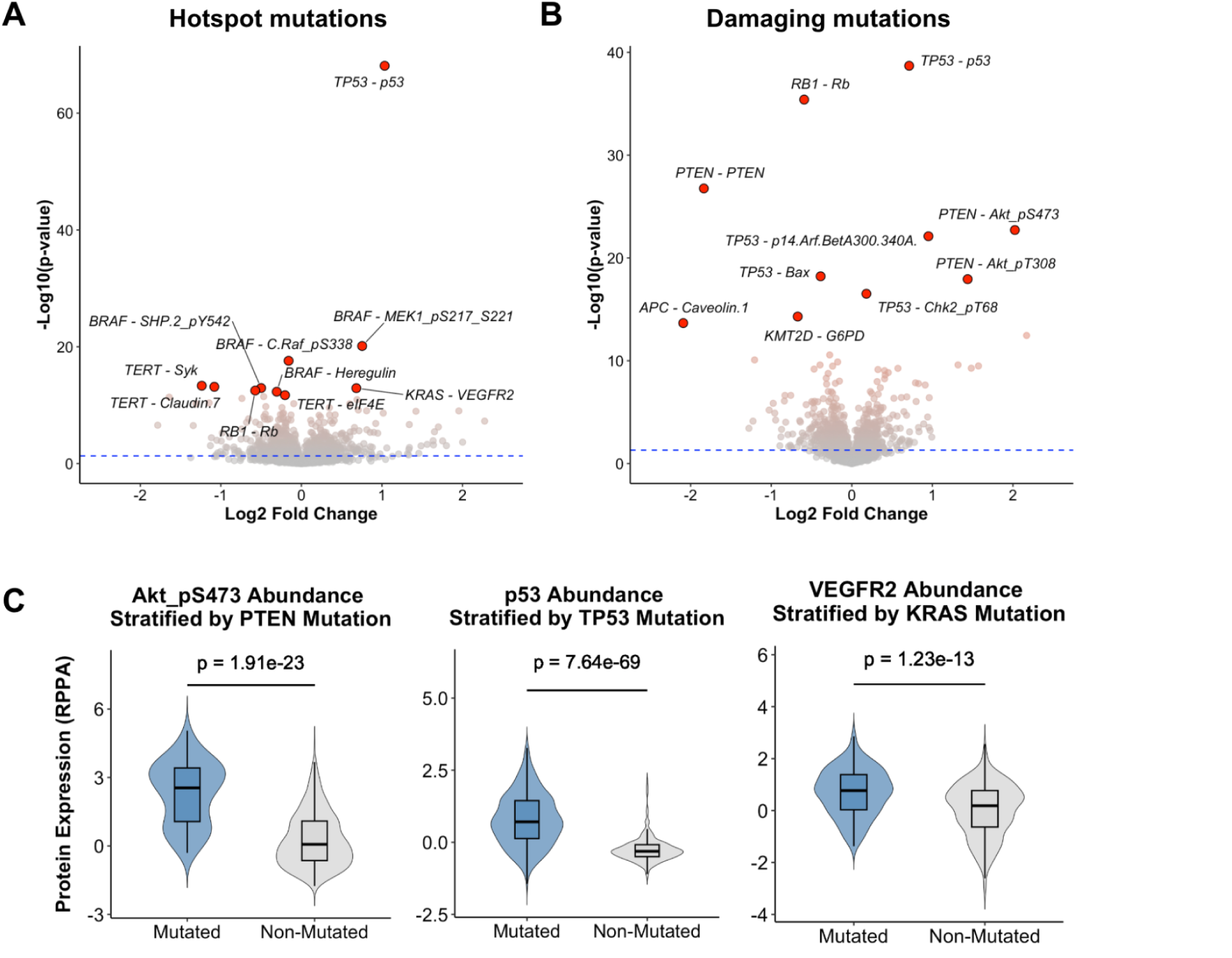
Gene and protein expression correlations with mutational status. A) Volcano plot showing top hotspot mutation - protein abundance relationships. Log2 Fold Change represents the enrichment in mutated lines relative to non-mutated cell lines. B) Volcano plot showing top damaging mutation - protein abundance relationships. Log2 Fold Change represents the enrichment in mutated lines relative to non-mutated cell lines. C) Violin plots visualizing select mutation - protein abundance relationships between Akt pS473 and *PTEN* mutation, p53 protein and *TP53* mutation, and VEGFR2 protein and *KRAS* mutation.

## Discussion

For many reasons (e.g. increased molecular profiling, AI and data driven approaches to molecular design, international efforts in China and elsewhere), there has been an explosion of new drug development in cancer. While high-effect-size drugs (e.g. chemotherapy) that were discovered over the last century by phenotypic screening still see widespread usage and have cured many patients, molecularly guided therapies – predominantly targeting genomic alterations such as mutations, amplifications, and copy number losses – have reshaped oncology treatment. However, this paradigm is reaching saturation, necessitating a more sophisticated understanding of the molecular contexts in which therapies are most effective. Enhancing biomarker strategies to better identify patients most likely to benefit from targeted agents can unlock both new and existing therapeutic targets.

Despite advances in targeted therapy, there has not been a corresponding expansion in biomarker diagnostics. A fundamental challenge remains: distinguishing responders from non-responders. Our analysis indicates that large cancer cell line repositories, such as DepMap, can serve as a valuable resource for identifying clinically relevant dependencies and their associated biomarkers for clinical translation. Our findings highlight several key observations with therapeutic significance:

- **High Prevalence of Paralog Dependencies:** Paralog dependencies where reduced expression of a paralog pair strongly enhances dependency on the other member (e.g. *VPS4A* dependency in cell lines with *VPS4B* loss^59^) are frequently observed. These dependencies can be readily detected using existing sequencing-based genotyping tests, such as Tempus xT and FoundationOne CDx which can detect copy number loss or loss of function mutations in paralog pairs, thus nominating targeting of the other paralog member.
- **SYDE1 as a Biomarker for Focal Adhesion Pathway Dependencies:** The focal adhesion pathway has been a long-standing target for therapeutic development^28^, but clinical trials have largely been disappointing due to low single-agent activity^60^. Our findings suggest that the expression of *SYDE1*, a Rho GTPase-activating protein encoding gene, is strongly correlated with dependency on multiple genes within this pathway. A deeper understanding of when cancer cells with high expression of *SYDE1* are most susceptible to focal adhesion inhibitors may help revive interest in this therapeutic avenue.
- **Mutation-Stratified PROTAC Targets:** Our analysis revealed that PROTAC (proteolysis-targeting chimera) targets can be identified through mutation stratification. While p53 remains a well-known example^55^, we also found that PTEN-mutated tumors accumulate Akt-phosphorylated proteins, a finding potentially translatable for certain induced proximity technologies. Although protein-based biomarkers would ideally be used for patient selection, mutation-based stratification could serve as an immediately implementable strategy in clinical trials.
- **Transcription Factor Dependencies as Selective Targets:** Our and others’ analyses highlight a striking opportunity to develop selective transcription factor targeting compounds for cancers dependent on aberrant usage of lineage associated TFs. Many TF dependencies are lineage-specific and strongly correlated with the expression of the oncogenic TF itself. However, caution is necessary—PAX8, for example, is critical for thyroid function but is also a key dependency in the reproductive system and kidney tumors. To fully evaluate the therapeutic potential of targeting TFs, it may be necessary to develop cell lines derived from normal tissues to assess the feasibility of selective targeting.
- **Underexplored Lineage-Specific Dependencies:** Our analysis also identified underexplored lineage-specific dependencies, which may offer specific areas for biological investigation to understand the pathogenesis of certain cancer lineages. For example, our approach identified the critical role of *ZER1* in cervical cancer, which contributes to the oncogenic activity of high-risk HPV E7 proteins^24^. Targeting these unique dependencies may open new therapeutic avenues in historically challenging cancers.
- **Expanding the Cancer Dependency Map:** A major limitation of current datasets is the underrepresentation of certain cancer types. For example, DepMap includes only 24 liver cancer cell lines, which may hinder the identification of lineage-specific dependencies. Beyond generating models from more patients, expanding cell lines to include post-therapy settings—such as neoadjuvant-treated tumors or those grown to resistance in increasing drug concentrations—could provide valuable insights into treatment resistance mechanisms. Currently, DepMap has not reached saturation in statistical power^61^, and further expansion could enhance the predictive accuracy of selective drug targets. Not all -omics measurements are strictly necessary. Our data suggests that RNA, DNA, and proteomic biomarkers appear to have the greatest statistical power for stratifying dependency and utility for functional analysis.
- **Proteomics as a Tool for Predicting Drug Sensitivity:** Our findings reinforce the value of expression measurements such as proteomics in distinguishing sensitivity to targeted therapies. Expression-based biomarkers—critical for predicting dependency—may be best detected using protein-based assays rather than RNA-based tests, which are more volatile and can be less interpretable. Additionally, unlike transcriptomic assays, proteomic approaches—such as immunohistochemistry (IHC)—are already integrated into clinical workflows. Expanding these techniques through high-plex proteomics (e.g., 40+ proteins per slide) could significantly enhance patient stratification. Protein-based biomarkers hold several advantages: high sensitivity, compatibility with single-cell analysis, routine clinical specimen use, multiplexing capability, and direct tumor material analysis (unlike liquid biopsy). However, large-scale implementation will require advanced computational resources for image analysis, stain quantification, and biomarker identification, as well as more specific and reliable antibodies. Beyond cancer, multiplexed biomarker analysis can be even more crucial, as many diseases lack a strong genomic driver. Measuring expression in patient-derived tissues (e.g., kidney biopsies for lupus nephritis, intestinal biopsies for inflammatory bowel diseases, and lung biopsies for fibrotic diseases) or fluids (e.g., blood, urine) may provide clinically actionable insights.

Our analyses were primarily constrained by database size and quality. The clinical viability of a given target depends not only on its dependency score but also on drug-specific properties such as off-target activity, potency, and clinical trial design, which can complicate a binary assessment of therapeutic potential. Additionally, many promising targets lack drugs in development, while others have yet to progress through clinical trials. Although CRISPR screening and gene expression datasets provided genome-wide quantification, our protein quantification panel was relatively limited, assaying only 214 distinct proteins and phosphosites. As the catalog of validated, cost-effective antibodies expands and unbiased protein measurement technologies—such as sequencing and mass spectrometry—become more accessible and standardized, the landscape of proteomic biomarker discovery will improve. However, we note that protein sequencing and mass spectrometry still face significant limitations in sensitivity, dynamic range, and specificity.

Looking ahead, the landscape of molecularly targeted therapy will likely become increasingly complex, making it more difficult to directly compare drugs while accounting for molecular and clinical heterogeneity. Beyond expanding the cancer cell line encyclopedia and developing new multiplexed tools for in situ protein biomarker measurement, establishing robust preclinical systems to predict therapeutic windows and facilitate head-to-head drug comparisons will be critical. The integration of large patient-derived model libraries multiplexed in situ biomarker assays that isolate biomarker signals specifically from cancer cells, and high-fidelity preclinical models represent a promising framework for refining cancer precision medicine. By advancing these tools and strategies, we can enhance patient stratification, optimize therapeutic targeting, and accelerate the translation of novel treatments into clinical practice.

## Methods

### Source of Data

Publicly available RNAseq, protein antibody array, copy number, mutation, PRISM drug screening, and CRISPR dependency data were downloaded from DepMap release Q42024. Methods for quantification of biomarkers are detailed by DepMap elsewhere^62^. In brief, gene expression levels were quantified as log_2_(TPM+1) and the gene copy number was computed relative to the ploidy of the rest of the genome for a given cell line. Mutations and chromosomal aberrations were identified using SNP array chips. Protein expression was quantified by Reverse Phase Protein Array (RPPA).

Dependency effect sizes are quantified as Chronos scores, a statistical measure inferred from a population dynamics model that corrects for sgRNA efficacy, screen quality and cell intrinsic growth rates, and bias related to DNA cutting toxicity^16^. Dependency scores are continuous values that range in magnitude but have empirically demonstrated that essential genes commonly have Chronos score < -1, and unexpressed genes commonly have Chronos score equal to 0. Data for clinically validated targeted therapy strategies was manually curated by a review of FDA approvals and current clinical trial data.

### Clinically Tractable Targets

Clinically tractable targets were defined via OpenTargets schema and retrieved on January 10th 2025. Targets with molecules in Phase I or beyond (ie. Phase II/III or FDA approved medicines) were included and defined as clinically tractable.

### Dependency Enrichment

Enriched dependencies were nominated by a two-sided t-test between gene dependency effect sizes in each lineage relative to all other cell lines. Genes with a negative *t*-statistic were considered to exhibit stronger dependencies in the lineage (*p* < 0.01). Dependencies were further evaluated by a skewed likelihood ratio test (LRT), and those that contained a skewed LRT metric of greater than 100 were identified as being ‘strongly selective.’ In brief, the LRT metric is defined as twice the log of the likelihood ratio of a fitted skewed distribution of gene sensitivity (across all lineages) over the likelihood of a fitted normal distribution^63^.

### Dependency Predictability and Biomarker Predictiveness

Biomarkers associated with dependencies were identified by Pearson correlation coefficients as well as the relative importance of the predictive feature, as detailed by DepMap^10^. In brief, the relative importance indicates the impact of the feature on prediction accuracy relative to other features in the model (mRNA expression, damaging mutations, driver mutations, hotspot mutations, lineage annotation, fusion, copy number, and confounder experimental covariates), is computed with Gini importance, and is normalized so that the sum of all feature importances add up to 100.

### GO Term Analysis

To analyze gene set terms associated with PDAC selective genes, we performed an over representation analysis using the g:Profiler web server (https://biit.cs.ut.ee/gprofiler/gost). Gene sets with less than 500 genes were included for analysis, and statistically significant signatures at an FDR-adjusted p-value of less than 0.01 were plotted.

### Data Analysis

All analyses were performed in R version 4.3.3. Unless otherwise specified, group comparisons were performed using an unpaired two-sided t-test. A cutoff of p < 0.01 was used for significance. All R code and any source data are available upon reasonable request.

## Figures

**Supplementary Figure 1:**
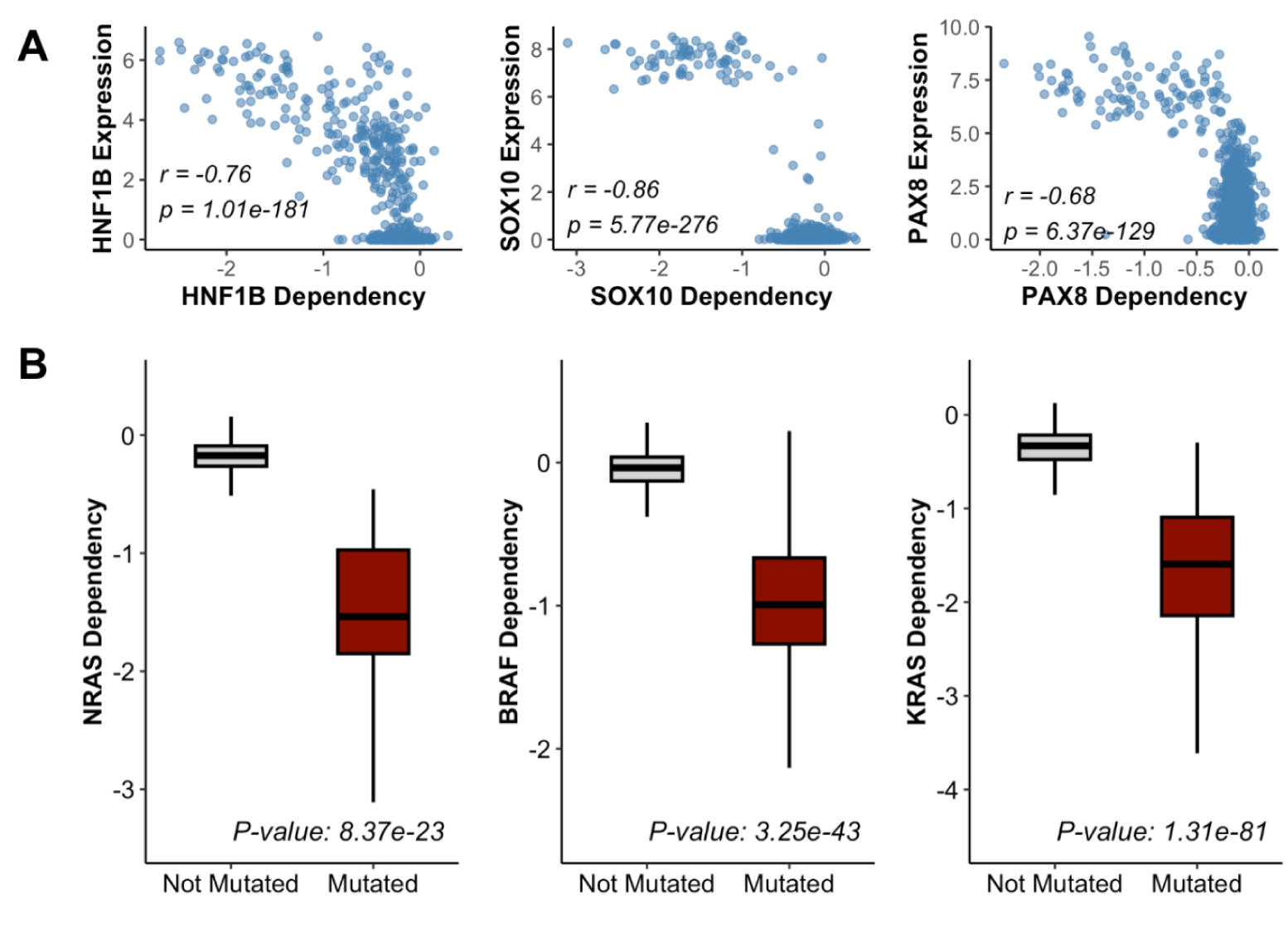
Visualization of strongest biomarker relationships. A) Scatterplot of *HNF1B*, *SOX10*, and *PAX8* RNA expression (y-axis) versus each gene’s respective dependency (x-axis). B) Boxplots comparing dependency of *KRAS*, *BRAF*, and *NRAS* in cell lines where the gene is mutated versus wild type.

**Supplementary Figure 2:**
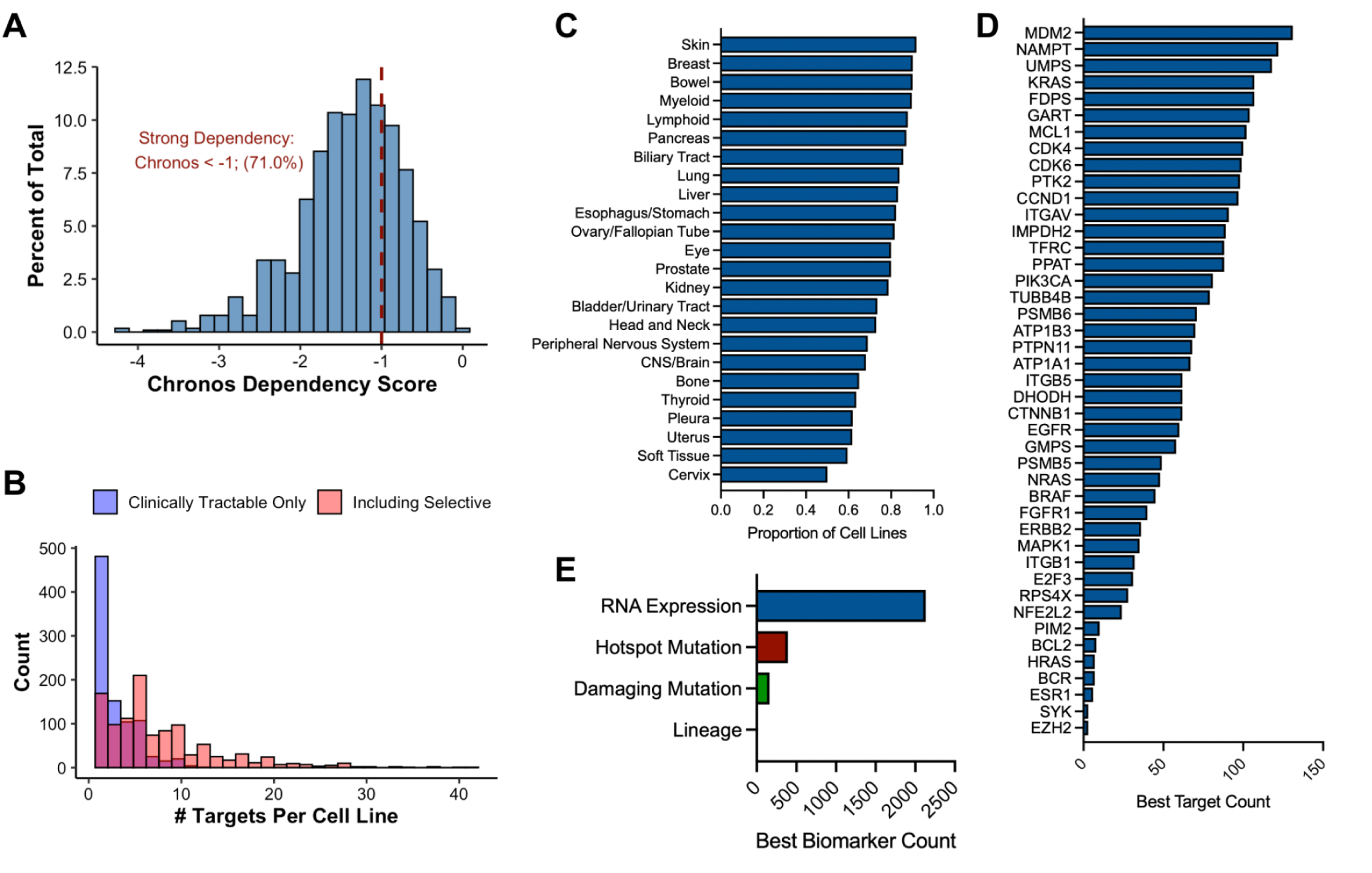
Nominated targets for each cell line. A) Histogram of Chronos dependency scores (x-axis) of the most preferential dependency for each cell line in DepMap. Dotted red line denotes threshold for strong dependency (Chronos < -1). B) Stacked histogram plotting the number of preferential targets (Chronos < 1.5 standard deviations below mean) per cell line with Chronos score < -1. Blue bars represent clinically tractable but predictable targets only. Red bars include selective targets. C) Barplot visualizing proportion of cell lines in each lineage that have preferential targets. D) Barplot visualizing most common preferential targets. E) Barplot visualizing most common biomarker classes for each preferential target.

**Supplementary Figure 3:**
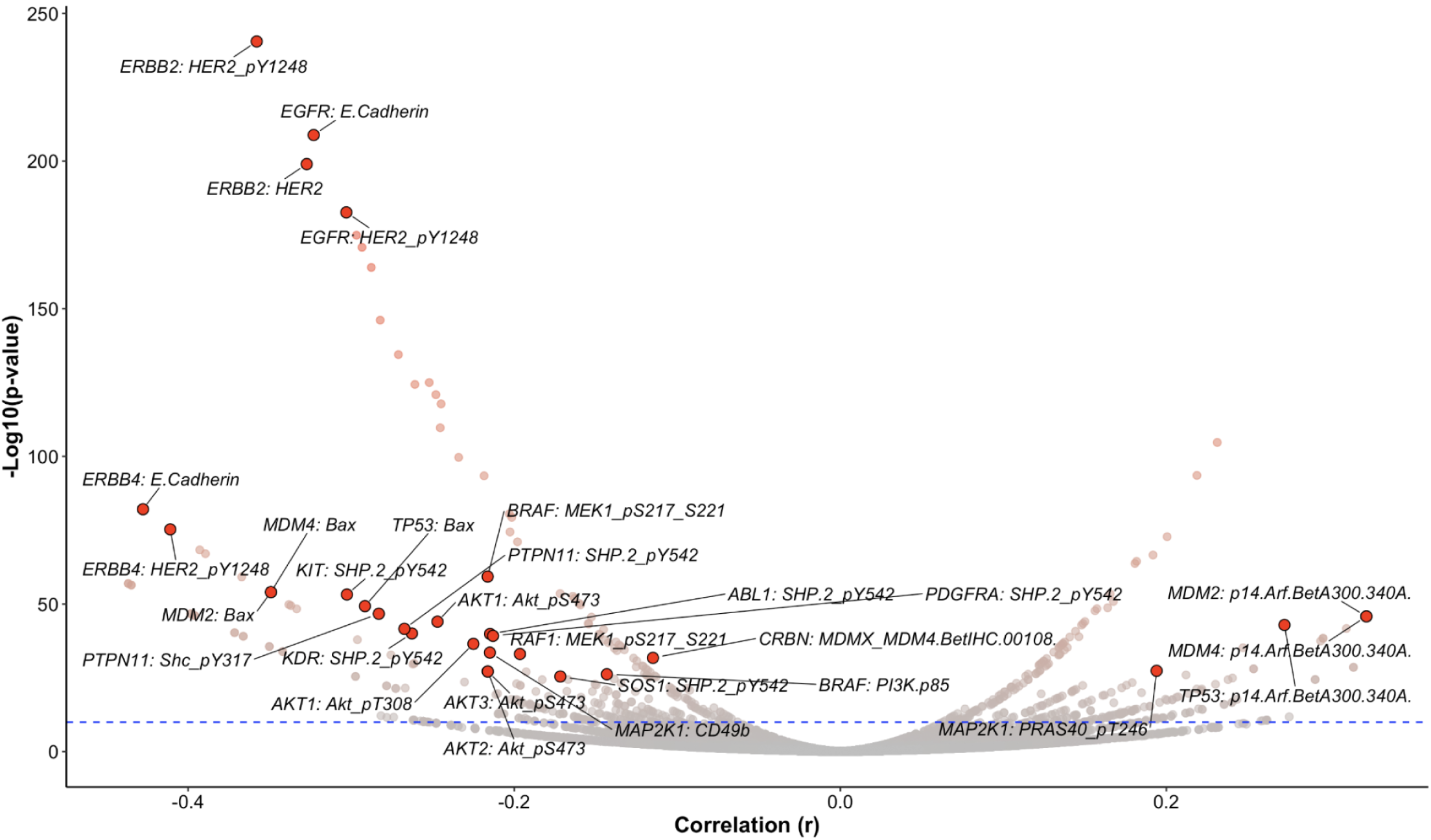
Protein biomarkers associated with dependency and small molecule sensitivity. Volcano plot showing relationships between a target’s aggregate PRISM drug screening sensitivity (based on all agents targeting the reported protein) and protein abundance. Correlation (r; x-axis) represents the correlation between PRISM AUC of drugs targeting the protein member, and protein abundance across all cell lines.

